# Machine learning prediction of antiviral-HPV protein interactions for anti-HPV pharmacotherapy

**DOI:** 10.1101/2021.08.22.457260

**Authors:** Hui-Heng Lin, Qian-Ru Zhang, Xiangjun Kong, Liuping Zhang, Yong Zhang, Yanyan Tang, Hongyan Xu

## Abstract

Persistent infection with high-risk types Human Papillomavirus could cause diseases including cervical cancers and oropharyngeal cancers. Nonetheless, so far there is no effective pharmacotherapy for treating the infection from high-risk HPV types, and hence it remains to be a severe threat to the health of female. Based on drug repositioning strategy, we trained and benchmarked multiple machine learning models so as to predict potential effective antiviral drugs for HPV infection in this work. Through optimizing models, measuring models’ predictive performance using 182 pairs of antiviral-target interaction dataset which were all approved by the United States Food and Drug Administration, and benchmarking different models’ predictive performance, we identified the optimized Support Vector Machine and K-Nearest Neighbor classifier with high precision score were the best two predictors (0.80 and 0.85 respectively) amongst classifiers of Support Vector Machine, Random forest, Adaboost, Naïve Bayes, K-Nearest Neighbors, and Logistic regression classifier. We applied these two predictors together and successfully predicted 57 pairs of antiviral-HPV protein interactions from 864 pairs of antiviral-HPV protein associations. Our work provided good drug candidates for anti-HPV drug discovery. So far as we know, we are the first one to conduct such HPV-oriented computational drug repositioning study.

## Introduction

Human Papillomavirus (HPV) can infect human body and cause different types of phenotypes. Specifically, HPVs can infect females’ reproductive system and causes different types of gynecological diseases. For instance, a variety of warts and genital cancers [1]. What’s more, it is reported that HPV infection is one of the risk factors of oropharyngeal cancer [2]. Researchers have classified the subtypes of HPVs into the low-risk types, and high-risk types according to their virulence and relevant risk levels of infections. For low-risk types [3], e.g., the type 6, 11, 40, etc, they might be disappeared after several periods of infection, and the infected hosts might generally be fine. While for those high-risk subtypes of HPV, e.g., the HPV-16 and HPV-18, their persistent infection on hosts could finally cause severe or lethal diseases like cervical cancer on hosts [4]. According to report, it is estimated that 569,000 cases of cervical cancer newly occurred in 2018 globally and 311,000 deaths were found [5]. Therefore, the HPV infection remains a large threat to female’s health especially in developing countries, and treating HPV infection remains an urgent task and difficult challenge due to there lacks effective pharmacotherapy. Though HPV vaccines are available, they are ineffective for those who have already been infected by HPVs [6].

Scientists have been trying hard to combat against HPV infection. For instance, several researchers have identified HPV’s E6 and E7 proteins to be the virulent tumorigenesis risk factors [7, 8], and parts of their molecular 3D structural conformations have been revealed through approaches of *in silico* simulation [9] and structural biology [10]. Other studies have tested, discussed, and reviewed the *in vitro* effects of existed drug, i.e., the Human Immunodeficiency Virus (HIV) protease inhibitor, on HPV proteins and cells infected with HPVs [11, 12, 13, 14, 15]. These reports targeting existed drugs for HPV treatments showed that, compared with *de novo* drug discovery, repositioning exited drugs is indeed the better and quicker strategy.

Nonetheless, drug efficacies from above evidences were moderate and no further progress is seen in later stages, e.g., in clinical contexts. And hence, above research progresses are yet far from being able to identify drug candidates with good therapeutic and anti-HPV potential. Limitation of them could be due to two reasons. One is that inappropriate compound or drug candidates have been chosen for testing. The other reason could be the number of drug candidates to be tested is too small. Testing only limited numbers of compound or drug candidates surely restricts the probability of identifying those appropriate ones.

In order to meet the urgent needs for effective anti-HPV drug discovery, based on target-oriented drug repositioning strategy, we collected and analyzed 96 antiviral drugs to do the relatively large-scale *in silico* screening for 9 HPV-16 proteins, so as to computationally and effectively identify effectively antivirals with good potential for targeting HPV proteins. Briefly, in this work, we constructed, benchmarked, and selected machine learning predictive models (also known as predictors) to predict antivirals that could have potential interactions with HPV proteins. This is because drug-target interactions are vital prerequisite of molecular therapeutic mechanisms. Through benchmarking, we selected the high-precision K-Nearest Neighbor (KNN) [16] and Support vector machine (SVM) [17] predictors to detect those confidence interaction pairs of antiviral-HPV protein.

To the best of our knowledge, no prior study similar to our work has been done. Lots of researchers predicted targets of drugs, compound-protein interactions, or protein-protein interactions using machine learning or other computational methods [18, 19, 20, 21]. However, so far as we know, no study has focused on studying relationships between antiviral drugs and HPV proteins.

## Methods

### Research question formulation

Theoretically, a therapeutic target and its drug molecule have interactive binding relation to each other. Therefore, trying to identify potential HPV protein targets of antivirals could be considered as a binary classification task, i.e., to predictively classify proteome of HPV into two classes of proteins. One class is HPV proteins which have potential interaction with drug molecules, and the other class is HPV proteins do not have potential interactions with drugs. Machine learning is state-of-art method to solve such binary classifications **(Fig. 1)**. Considered that known antiviral drug-target interaction pairs were available, which could serve as the known-label validation dataset, we thus chose supervised (machine) learning methods for this study.

**Figure 1:**
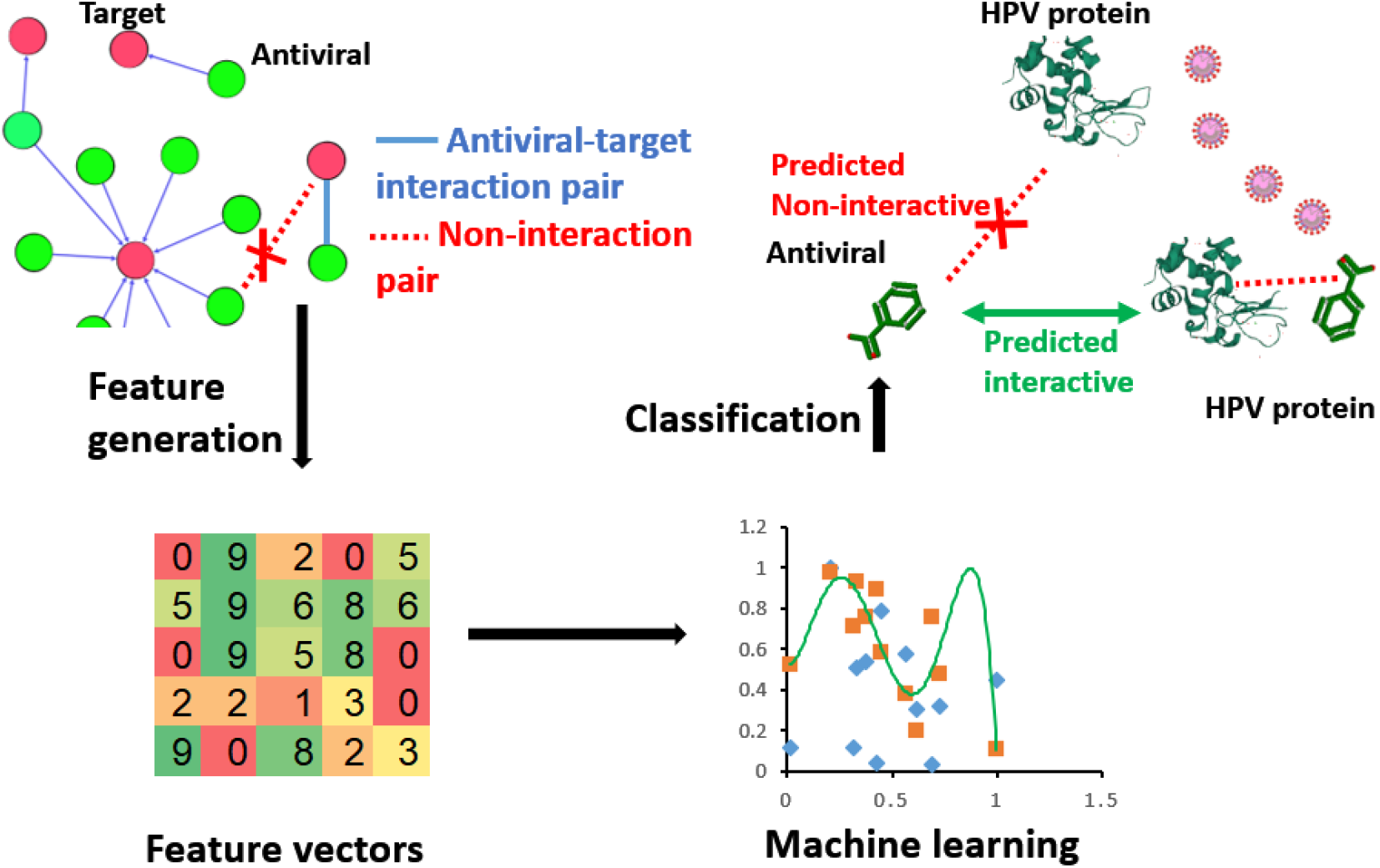
Research framework of this study. Predicting antiviral drug-HPV protein interaction could be considered a binary classification task, and machine learning is a good method for such task. In this work, antiviral drug-target pairs’ features were transformed into vectors for constructing machine learning predictors. Through benchmarking, the best predictors were selected to predict antiviral-HPV protein interactions.

### Data collection and preprocessing

We collected antiviral drugs and their associated target data from DrugBank [22], Drugs@FDA [23], PubChem [24], Uniprot [25] and Therapeutic Targets [26] databases (As of 19th July 2020). Drug-target interaction pairs which contained the United States Food and Drug Administration (FDA)-approved antiviral drugs were treated as the validation dataset for machine learning, because FDA-approved antivirals as the validation set can better reflect the real-world application value of our models. And the rest drug-target interaction pairs were treated as the training dataset for machine learning. In this work’s machine learning classification task, an interaction pair of an antiviral drug and a protein was defined to be a positive instance, while negative instance indicated a non-interactive pair of antiviral and protein. In order to balance data ratio for binary machine learning classification task. We randomly generated non-interactive drug-target pairs so as to assure the 1:1 ratio of positive instances to negative instances for machine learning. In more details, we initially constructed a full graph of bipartite drug-target network, in which each antiviral was connected to all the target proteins in the network. Upon removing those known antiviral-target interaction pairs, we had those non-interactive drug-target pairs. And then, we randomly drew such non-interactive drug-target pair out without replacement (treated it as the negative instances for machine learning) until the ratio of positive to negative instance reached 1:1. Next, we integrated the proteome (9 proteins in total) of high-risk HPV-16 subtype and all the antiviral drugs to form drug-protein interaction prediction dataset. See **Supplementary Table 1** for machine learning training dataset of antiviral drug-target interaction pairs, **Supplementary Table 2** for drug-target interaction pair dataset used in machine learning validation process, and **Supplementary Table 3** for Uniprot’s HPV-16 proteome, i.e., 9 proteins.

Next, all antivirals’ molecular structures were analyzed using ChemmineR [27] and 1024-dimension chemical fingerprint datasets were generated through R scripting [28]. All proteins were analyzed using ProtR [29] and 10784 high dimension protein descriptor feature datasets were generated. As seen in **Table 1**. Descriptors used were protein structural and physicochemical properties. These descriptors have been widely used in studying protein-protein interactions and protein-ligand interactions *in silico,* and they worked well [29]. All datasets were integrated, scaled and normalized using R computing environment [28].

**Table 1:** Molecular descriptors used for machine learning analysis

| ID | Molecular descriptor | Vector length |
| --- | --- | --- |
| 1 | Drug molecule fingerprint | 1024 |
| 2 | Amino acid composition | 20 |
| 3 | Dipeptide composition | 400 |
| 4 | Tripeptide composition | 8000 |
| 5 | Normalized Moreau-Broto Autocorrelation | 240 |
| 6 | Moran Autocorrelation | 240 |
| 7 | Geary Autocorrelation | 240 |
| 8 | Composition descriptor | 21 |
| 9 | Transition descriptor | 21 |
| 10 | Distribution descriptor | 105 |
| 11 | Pseudo amino acid composition | 50 |
| 12 | Amphiphilic pseudo amino acid composition | 80 |
| 13 | Conjoint Triad | 343 |
|  | Total | 10784 |

### Machine learning and prediction

Briefly, the machine learning processes of this research work followed such order and general steps. Initially, the training dataset was loaded to different machine learning algorithms, and 5-fold cross validation and grid searching were applied to training processes, so as to identify the best parameters of machine learning models with the best predictive power. Later, predictors with good performances were further applied to classify the validation dataset with known labels. Lastly, the verified best predictor was used to predict antiviral-HPV protein interaction pairs.

Diverse sorts of supervised learning algorithms with different purposes exist. Amongst, the Support Vector Machine, Random Forest [30], Logistic Regression [31], etc, are classic algorithms for tackling the binary classification questions. 6 types of machine learning classifiers friendly for binary classification were chosen for building predictive models. The chosen predictors were Support Vector Machine, Random Forest, AdaBoost [32], Logistic Regression, Naïve Bayes [33] and K-Nearest Neighbor classifier. Amongst, K-Nearest Neighbor classifier and Adaboost displayed good prediction performances on predicting miRNA-disease associations [34, 35]. And Chen et al. developed a Random Forest-based model RFMDA which had good predictive power on multiple kinds of human complex diseases [36]. These studies support us to choose aforementioned predictors for this work.

With default parameters, 6 predictors went over simple checking through quick training and performance measurement. At this early stage, as expected, all predictors did not perform well. Subsequently, in order to identify better parameters for predictors, grid search 5-fold cross validations and performance benchmarking were conducted. The predictive performance of 6 different predictors with better parameters were tested using known-label validation dataset. Upon checking performance of different predictors, we selected the optimized K-Nearest Neighbors classifier and SVM, which had the highest precision scores and were the most appropriate predictor to identify high confidence drug-protein interaction pairs from 864 pairs of antiviral-HPV protein associations.

Aforementioned data processing and machine learning computations were done via in-house scripts of Python [37] and R [28]. Libraries and modules used were Sci-kit learn [38], Pandas [39], Numpy [40, 41] Scipy [42], and also Bioconductor [43, 44] and Biomart [45].We acknowledge the authors and developers of these computational tools.

Specifically, parameter set of KNN that finally used for predicting antiviral-HPV protein interaction pair was that, the number of neighbors was set to 65, “weights” was set to “distance”, and “leaf_size” was set to 60. And for SVM, gamma was set to 0.001, C (the regularization parameter) was set to 0.0002, and polynomial kernel with degree = 3 was used to predict antiviral-HPV protein interaction pair. The rest parameters remained default ones of the function of Python library Sci-kit learn [38].

## Results

### Dataset overview

The antiviral drugs and their associated targets were retrieved and analyzed as described in method section. **Table 2** provides a summary of our dataset. We had totally 61 antiviral drugs, which formed totally 284 antiviral-target interaction pairs with their targets. For the purpose of measuring machine learning predictors’ performance, antiviral-target interactions were split into two classes where 102 pairs were used as the dataset for training or fitting machine learning predictors, and the rest 182 pairs were treated as dataset for validating the predictive performances of machine learning predictors. And we also compiled 9 proteins of HPV (its complete proteome) with 96 antiviral drugs to form 864 pairs of antiviral-HPV protein association pairs (**Table 2**).

**Table 2:** Summary of antiviral-target and antiviral-HPV protein interaction dataset used in machine learning processing of this study

| Dataset | Antiviral drug | Target protein | Antiviral-protein interaction pair |
| --- | --- | --- | --- |
| Training set | 35 | 34 | 102 <sup>c</sup> |
| Validation set <sup>a</sup> | 61 | 47 | 182 <sup>c, d</sup> |
| Prediction set | 96 | 9 <sup>b</sup> | 864 |

<sup>a</sup> Validation set consisted of U.S.FDA-approved antiviral drugs and these drugs’ binding target proteins
<sup>b</sup> 9 proteins of HPV-16.
<sup>c</sup> Ratio of positive instance to negative instance was 1:1.
<sup>d</sup> Number of validation set was greater than that of training set because (1) more FDA-approved antivirals were desired for validating the real-world application value of our machine learning models; (2) generalization performance of machine learning models could be reflected using smaller training set but larger validation set.

### Performances of machine learning models

Initially, we chose 6 types of machine learning models and applied 5-fold cross validation strategy to fit the antiviral-target interaction training dataset. A primary benchmarking of the predictive performance of 6 chosen predictors was as seen in **Table 3**.

**Table 3:** Performance of 6 machine learning predictors with default parameters.

| ID | Predictor | Precision | Recall | F1-measure | Accuracy | AUC <sup>a</sup> |
| --- | --- | --- | --- | --- | --- | --- |
| 1 | SVM | 0.36 | 0.48 | 0.37 | 0.48 | 0.44 |
| 2 | Logistic Regression | 0.52 | 0.57 | 0.54 | 0.59 | 0.56 |
| 3 | KNN | 0.61 | 0.62 | 0.60 | 0.59 | 0.61 |
| 4 | Naïve Bayes | 0.46 | 0.65 | 0.52 | 0.49 | 0.52 |
| 5 | Random Forest | 0.61 | 0.61 | 0.63 | 0.66 | 0.68 |
| 6 | AdaBoost | 0.73 | 0.48 | 0.55 | 0.62 | 0.48 |

<sup>a</sup> AUC indicates the metric of Area Under Curve of Receiver-Operating Characteristic Curve.

Briefly, all predictors’ predictive performances were less satisfying, as expected. SVM with default parameter (RBF kernel) performed the worst in all sorts of metrics among 6 predictors. AdaBoost classifier scored the best in terms of precision score but had the lowest recall score. F1-measure is the harmonic average of precision and recall. The highest F1-measure was found from the Random forest classifier, which was 0.63. While we also found other metrics of Random forest were not high. All its metrics were around 0.65 though the values were close to each other. The highest accuracy score and AUC (Area Under Curve of Receiver-Operating Characteristic Curve) of 6 predictors’ were 0.66 and 0.68, respectively. And both of them were also found in Random forest’s performance. Metrics of default parameters’ KNN were all around 0.6, indicating its unsatisfying performances in 5-fold cross validation, too. Similar to KNN, Naïve Bayes classifier did not perform well, and one common point of KNN and Naïve Bayes classifier was that, the value of their recall score was higher than those of other metrics (**Table 3**).

Next, we tuned parameters of predictors through grid searching 5-fold cross validation, and tested how combination set of parameters affected predictors’ predictive performances on known-label validation dataset. At the beginning, we focused on optimizing predictors for obtaining better values of comprehensive metrics, such as the F1-measure, accuracy or AUC value. Despite a great number of times’ trying, no high sores of aforementioned F1-measure, accuracy or AUC metric value was seen.

Given that high precision score indicates the low number of predictive false positive instances, and high recall score indicates the low number of predictive false negative instances, we changed our strategy and decided to do high precision-oriented optimization. This was because the purpose of this work was to identify antivirals that interact with HPV proteins. To this end, using high-precision predictor, predictive positive instances could have lower false positive instances mixed inside. Therefore, in this work, we preferred precision metric over recall metric for selecting appropriate predictors to predict antiviral-HPV protein interactions (positive instances). Through benchmarking the performances of predictors, we found optimized SVM and KNN predictors had better precision scores than others. SVM’s was 0.8 and the KNN classifier’s was 0.85 **(Table 4)**. We hence used them for prediction task and we chose the intersection of their prediction results as the final results.

**Table 4:** Precision scores of optimized machine learning predictors on the validation dataset of antiviral-HPV protein interaction

| Predictor | SVM | Logistic Regression | KNN | Naïve Bayes | Random Forest | AdaBoost |
| --- | --- | --- | --- | --- | --- | --- |
| Precision Score | **0.80<sup>a</sup>** | 0.50 | **0.85<sup>a</sup>** | 0.65 | 0.68 | 0.75 |

<sup>a</sup> Metrics of optimized SVM and KNN used for predicting antiviral-HPV protein interaction are available at Supplementary Table S4.

### Predicted antiviral-HPV protein interaction pairs

Upon selection of high-precision predictors, we applied them to predict the antiviral-HPV protein interactions. We selected two predictors’ result intersection as the final prediction result, i.e., we only consider an antiviral-HPV protein association pair has potential interaction if both predictors predicted this pair to be position (interactive). As a result, within 864 antiviral-HPV protein association pairs, most antiviral-HPV protein pairs were predicted to be negative, i.e., the antiviral drug does not interact with the HPV protein. Only a small portion, i.e., 57 of antiviral-HPV protein pairs were predicted to have interaction. Prediction results were summarized in **Table 5** in HPV protein-oriented form. Full prediction results could be found in **Supplementary Table 3**. Here we took the Docosanol as an example for analysis. The drug Docosanol was predicted to interact with HPV-16’s protein E7 using our high-precision machine learning predictors. Docosanol is a U.S. FDA-approved antiviral drug targeting Envelope glycoprotein GP350 and GP340 of Epstein-Barr Virus (EBV, also known as Human Herpesvirus or HHV-4) and it is used to treat fever blisters, etc. Interestingly, through literature survey, a recently published clinical case report was found to claim that, the mixture usage of Docosanol, curcumin, and other drugs together treated HPV infection and vaginal warts of a patient well [46]. This could be evidence supporting our predictive result about Docosanol and HPV protein. HPV protein-oriented antiviral prediction results were summarized in **Table 5** and brief description of the example antiviral drugs, protein targets of the antiviral drug and relevant therapeutic indications were also listed in **Table 5**.

**Table 5:** Summary of prediction result of antivirals targeting each protein of HPV-16.

| HPV-16 protein | Number of antivirals <sup>a</sup> | Example |
| --- | --- | --- |
| Protein E7 | 7 | Docosanol targeting GP340 or GP350 protein of Epstein-Barr Virus has been approved to treat herpes labialis, fever blisters, etc. |
| Regular Protein E2 | 5 | Voxilaprevir targeting NS3/4A protein of Hepatitis C Virus has been approved to treat chronic Hepatitis C caused by Hepatitis C Virus infection. |
| Protein E6 | 6 | Telaprevir is an NS3/4A viral protease inhibitor. It has been approved to treat chronic Hepatitis C Virus infection in combination with other drugs. |
| Minor capsid protein L2 | 4 | Grazoprevir targeting NS3/4A protein of Hepatitis C Virus has been approved to treat Hepatitis C viral infection. |
| Protein E4 | 8 | Nelfinavir is a potent viral protease inhibitor for treating infections of Human Immunodeficiency Virus (HIV), and it targets the protease of HIV -1. |
| Probable protein E5 | 7 | Maraviroc is a chemokine receptor antagonist drug targeting C-C chemokine receptor type 5. It has been approved to treat HIV-1 infection |
| Replication protein E1 | 7 | Pirodavir (investigational drug) targets the genome polyprotein of Polioviruses and it seems to have broad-spectrum antiviral effects on multiple kinds of Human Rhinoviruses. |
| Major capsid protein L1 | 5 | Docosanol targeting GP340 or GP350 protein of Epstein-Barr Virus has been approved to treat herpes labialis, fever blisters, etc. |
| Protein E8^E2C | 7 | TMC-310911 (investigational drug) is a protease inhibitor targeting HIV-1 protease and it seems to have effect on treating HIV-1 infection. |

<sup>a</sup> Indicating the number of antivirals which was predicted to have potential interaction with specific HPV-16 protein.

## Discussion

While our results are to be validated by *in vitro* assays, in this work, we constructed machine learning models, and predicted antiviral-HPV protein interactions so as to identify potential drug candidates targeting HPV proteins. The high-risk types of HPV are not limited to HPV-16. There are other types such as HPV-18. Indeed, we are not only able to apply the research framework of this study to predict the potential drug candidates for the proteome of other HPV subtypes, but also to other types of pathogenic and infectious microbes, as well.

Reviewing this current study, we found several significant points that could help us do better preparation for further works. Initially, in this work, though we tried our best to collect more antiviral drugs, due to the availability of antiviral drugs, we had limited size of dataset for machine learning. This could be one of factors why we did not obtain predictors with high scores of F1-measure, accuracy, or AUC. Compared with antivirals, the amount of other types of drugs, e.g., cancer drugs or antibiotics, is higher. Thus, in future studies, we would consider using other types of drugs for repositioning purpose.

Also, the final predictors selected did not have high F1-measure, accuracy, or AUC. Because current machine learning processes are black box which is difficult to interpret. Alternatively, in this study, considered the tradeoff between precision and recall, we chose to select the intersected prediction results from two high-precision predictors in order to get higher confidence antiviral-HPV protein interactions. For future study, we would learn and try to apply the state-of-art explainable machine learning methods which may be interpretable. In such case, we may be able to find out reasons causing low performances and obtaining guidance for model optimizations and obtaining more powerful machine learning predictors. One more interesting idea for extending current work is to predict synergistic antiviral drug combinations for HPV infection pharmacotherapy. Similar to “cocktail” treatment for HIV infections and synergistic treatment for fungal infections, it is likely that synergistic drug combinations work for treating HPV infections, too. A good example to get insights from is NLLSS [47], which is a well-performed algorithm for predicting antifungal synergistic drug combinations. Similarly, it is a computational and machine learning-based research work, and hence multiple points, such as its research ideas and methodology, could be referred to.

## Conclusions

Inspired by the needs of anti-HPV drug discovery, drug repositioning and computational analytics, we designed this research project and constructed machine learning models to predict possible antiviral-HPV protein interactions so as to identify potential pharmacotherapy for HPV infection.As a result, we optimized the predictors and identified 57 antiviral-HPV protein interaction pairs.

To the best of our knowledge, we are the first pioneer to conduct this HPV-oriented computational antiviral repositioning study. No similar study has been found so far. Therefore, our work provides good insights to virologists, medicinal chemists, gynecologists, clinical microbiologists, etc, those who are interested in the treatment and therapy of HPV infections. Also, drug candidates pre-selected via computational analytic screening could have lower probability of ineffectiveness than those that did not go through computational analyses. It thus could save resources, and antivirals identified by us could be good candidates for further *in vitro* and *in vivo* tests. In such way, this work contributes to drug development for HPV infections. What is more, our predicted antiviral-HPV protein interaction pairs also offer insights for fundamental biomedical research on drug-protein interactions or molecular interaction mechanisms. The last but not the least, the research framework of this study, i.e., the machine learning-based compound-protein interaction prediction, could also be applied to primary drug repositioning or drug discovery for those diseases or infectious microbial pathogens lacking effective pharmacotherapy. E.g., the Noroviurs and COVID-19.

## Author Contributions

- Study conceptualization and design: Hui-Heng Lin and Hongyan Xu;
- Funding acquisition: Hui-Heng Lin;
- Resources: Hui-Heng Lin;
- Methodology: Hui-Heng Lin;
- Project administration: Hui-Heng Lin;
- Supervision: Hui-Heng Lin;
- Data collections: Hui-Heng Lin, Xiangjun Kong and Qian-Ru Zhang;
- Data curation: Hui-Heng Lin;
- Investigation: Hui-Heng Lin;
- Data and formal analyses: Hui-Heng Lin and Liuping Zhang;
- Software/Programming: Hui-Heng Lin;
- Validation: Hui-Heng Lin;
- Writing orginal draft of manuscript: Hui-Heng Lin;
- Manuscript proofreading: All authors;
- Manuscript editing and revisions: Hui-Heng Lin, Yong Zhang and Yanyan Tang;
- Approval of submission: All authors approved submission.
- Visualization: Hui-Heng Lin;

## Completing Interests

Authors declare no competing interest.

## Data Availability Statement

Data of this study were included in the supplementary materials

